# Treatment and reprocessing of textile dyeing wastewater by using the electrocoagulation treatment process

**DOI:** 10.1101/2024.11.26.625502

**Authors:** Noor Fatima, Irfan Ahmad Shaikh

**Affiliations:** College of Earth and Environmental Sciences University of the Punjab 54590, Lahore, Pakistan

**Keywords:** reactive dyes, wastewater treatment, textile industry

## Abstract

This study electrocoagulated textile effluent to remove dyes. The wastewater reuse method was used to study three reactive dyes: C.I. Reactive Red 221, Blue 19, and Yellow 145. Laboratory wastewater was electrocoagulated. Standard wastewater had three pH values. (4, 7, 10). Electrocoagulation removes volatile dye colors well. The best results were 98% within 15-20 minutes for 1% C.I. Reactive Red 221. After 20 minutes, 1% shade C.I. Reactive Blue 19 and Yellow 145 color removal efficacy is 96%. Electrocoagulation treatment of dyeing effluent is greatly affected by pH. Electrocoagulation efficacy was lowest at pH 4. When pH was highest, removal efficacy was best. Dyeing wastewater removal efficiency improved when pH was changed from 7 to 10. Discolored wash-off cloth batches were rubbed dry and wet. The normal and electro-coagulant treated wash-off batches had wash fastness and rubbing values of 4 to 5 in dry crocking. The C.I. Reactive Dyes (Red 221, Yellow 145, and Blue 19) standard fabrics and treated wash-off with electro-coagulated lot showed 3 to 4 wet rubbing, almost like the standard fabric, but no shade changes at this concentration. All three reactive dyes had color values. Because alkaline pH removes dyes best, color difference values of Batch C (fabric treated by pH 10 wash-off) were within the approved range of 0.38 to 1.50 of all dyes and shades (5%,3%,1%). Batch A values between 2.44 and 13.48 were outside the allowed range (< 1). Batch B (fabric treated by pH 7 wash-off) values fell between Batch A and Batch C.

## 1. Introduction

Water shortage is one of the biggest challenges that humankind faces on this planet Earth. This lack of water is increasing daily, and due to the unwise use of water and its overconsumption by the human population, it is continually growing, and our survival on earth becomes doubtful. (Kumar et al., 2022)Various water treatment techniques (which techniques) have been introduced to overcome this issue. These technologies are highly efficient, cost-effective, and reliable, so the wastewater generated on a large scale can be treated and reused. However, some of these technologies require a large amount of energy, which becomes a source of carbon emissions, and hence, these technologies are not environmentally friendly (Hossain et al., 2013). Water is an essential natural source in maintaining economic development and ecological balance. Due to progress in industrial development and more pressure on water resources, water consumption has tremendously increased, leading to water shortages (Bayramoglu et al., 2007).

The textile industry has an essential position around the globe. Because they contribute a significant role in the development of the economy and in fulfilling human desires according to their demand. Textile industries are the prime contributors to using dyes, water, and other toxic compounds. These liquids are generated from various processing units and contain various useable and unusable substances, which have serious environmental consequences. Due to these poisonous substances, the textile industry waste should be treated before entering the environment. Treating this toxic water before its discharge into the environment saves the environment from various harmful effects (Dasgupta et al., 2015).In Pakistan, the textile industry is considered the largest manufacturing industry, with 2.5 crore employees. Pakistan had six spinning mills in 1947, but they have increased to 503. The main factor for growth is the decrease in tariffs. From 1985, the cotton import rate decreased from 85-50 %, and in 1988, further reduced by 20%. After 1988, they removed the restriction on expanding textile industries, so their number increased rapidly. Pakistan has become the world’s fourth-largest cotton producer, second only to China, the US, and India. The textile industry is essential for strengthening Pakistan’s economy and earning foreign exchange. In 1 year, it accounts for 60-65% of earnings in foreign exchange (Mansor et al., 2020). The industry consists of large, organized sectors and highly dispersed sub-units. Spinning is the central part of the value of the various textile industries. Many spinning industries have their operations organized in a way that weaves internally before and after the dyeing treatment facilities(Burkinshaw & Salihu, 2013). The processing department includes the printing, dyeing, and post-press processing sub-sectors. They operate mainly as small and medium-sized units(Kobya et al., 2003).

The printing section dominates all the processes in the textile industry. The dyeing and bleaching of the fabric follow the printing section. In the textile value chain, garment manufacturing creates the most significant employment opportunities. Small division units account for over 75% of all other divisions (Shah et al., 2014). Most knitwear industries operate as a general unit, i.e., processing, makeup facilities, and knitting. Big cities of Pakistan like Karachi, Lahore, and Faisalabad have a high concentration of the textile and knitwear industries where they have enough female labor. Pakistan is the third-biggest consumer of cotton and the fourth-largest producer globally (Chantes et al., 2015).

In the past 50 years, the textile industry has been Pakistan’s main driving force for foreign exchange earnings and job creation. The garment and textile industry is a vital engine that will increase the economy in the future (Verma, 2017). No other industry or service sector benefits the economy through its foreign exchange earnings and the creation of new employment opportunities, especially if there is synergy between different sub-sectors and efforts are made to develop actively (Shah et al., 2014).

### Textile Industry Process

Various technical procedures for machinery are vital in forming the required shapes and final products. Many effluents are released during multiple steps such as sizing, de-sizing, bleaching, scouring, and mercerization (Li et al., 2021). They are also generated during dyeing, printing, and finishing. The composition of such effluents mainly comprises dye residues, salts, acids, bases, various chemical agents, and byproducts. Multiple salts, such as sodium chloride (NaCl) and sodium sulphate (Na2SO4), were obtained as a byproduct of neutralization during wet processing in the textile industry (Verma, 2017).

### Methods used for Wastewater Treatment

There are various methods for the treatment of wastewater. These treatments have been divided into 3 main categories commonly named physical, chemical, and biological methods. Further processes are involved in these methods. Many conventional methods are also used to treat textile wastewater. These methods include the physical, physiochemical, or biological.

Because textile wastewater is diverse in composition, traditional methods (Heebner & Abbassi, 2022) are not sufficient or adequate to remove all organic or inorganic material from the water. It becomes compulsory for every industry to release its effluent in a treated form that meets the quality of specified standards (Garcia-Segura et al., 2017).

### Electrocoagulation Process

Electrocoagulation treats all solids (suspended, colloidal, and dissolved), metals, and dyes from the wastewater. The electrocoagulation process also helps remove pesticides, pollutants, and harmful microorganisms from the water. In EC operation, two electrodes of iron or aluminum are dispersed to treat water. The power supply source is DC (Zazou et al., 2019). When the EC process runs, the metal ions from the electrode dissolve. The metal hydroxide produced at a suitable pH is insoluble in water and can be removed. There is no further addition of chemicals; thus, no secondary pollutant is produced. EC method uses simple equipment, and sludge production is minimal.

## 2. Materials and Methods

**Lab Scale Set Up** Steps involved in the lab–scale setup include:

### Materials

Fabric, chemicals, and dyes are used in this study.

#### Fabric

4 samples of 5 g of cotton fabric were used for dyeing separately. From that, 1 batch was used as standard, and the remaining 3 were Batch samples to treat them at different pH (4, 7, 10). Three different dye shades were used (1%, 3%, 5%) for each reactive dye at a liquor ratio (L: R) 1:10. Isothermal dyeing process for each type of dye at 60°C for 30 minutes was accomplished as per standard technique. After dyeing, each sample was passed through a wash-off process.

#### Dyes

Different synthetic reactive dyes were used: C.I. Reactive Red 221, C.I. Reactive Blue 19, and C.I. Reactive Yellow 145 dye. The properties of dyes are given in table 1,2 and 3.

**Table 1.** Wash off the treatment process.

| **Wash-off of Batch Fabrics** |  |  |
| --- | --- | --- |
| **Fabric** | **pH** | **Wash-off Process** |
| **Standard** | 6.3 | Conventional Method |
| **Batch A** | 4 | Washed-off by treating the standard wash-off wastewater using the Electrocoagulation technique. |
| **Batch B** | 7 |  |
| **Batch C** | 10 |  |

#### Chemicals

Chemicals used include salt, acid, base, and soaping agents. Sodium Chloride (NaCl) and Sodium Carbonate (Na2CO3) were used in the dyeing process as catalysts for dye fixation on fabric. Tap water was used to prepare and dye wastewater. During the wash-off process, acid (Acetic acid) and alkali (Sodium hydroxide) were used to neutralize water (Khandegar & Saroha, 2013). A few drops of Soaping agent were added to the water during the soaping wash-off process.

**Figure 1.**
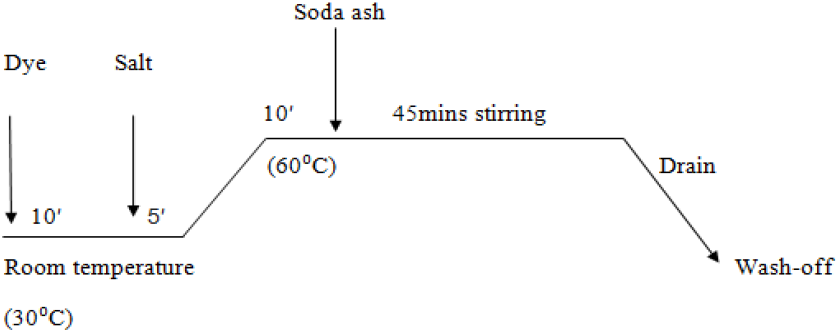
Standard Isothermal Dyeing

### Instruments

The instruments of concern were the Thermostatic Water Bath, pH meter, Spectrophotometer, weighing balance, thermometer, and Filter papers.

### Electro coagulator

The electrocoagulator with two electrodes was used to treat dyeing wastewater. The electrodes were made of iron, and the distance between them was 2.5 cm (Kumar et al., 2022). The electrodes were connected parallel to each other, and the current supply source was DC. Due to its process design, the EC process is preferred over physiochemical or biological methods. The material used in EC treatment is cost-effective.

To protect the environment, organic and inorganic materials are removed from wastewater. Textile wastewater with a specific dye composition and different pH analyses was done. Three reactive dyes, red 221, blue 19, and yellow 145, at pH 4, 7, and 10, were used after the whole analysis. It was found that in a high alkaline medium, which is at pH 10, the electrocoagulation technique gives the best results.

### Methodology

Standard and Batch Fabrics were dyed using an isothermal process, which used alkali, salts, and other chemicals. A liquid ratio of 1:10 was maintained for the dyeing and wash-off processes. The wash-off wastewater was treated using Electrocoagulation (Amour et al., 2016).

### Dyeing Method

A conventional dyeing process was used to dye standards and batch samples. The same dyeing method was used for 1%, 3%, and 5% depth shades, but the amount of Reactive dye, salt, and alkali varied for each depth shade.

#### 5% Dyeing Depth Shade

For 5% depth shade dyeing, 50 ml water, 0.25g Reactive dye, and 4g/50 ml NaCl were taken in a beaker. Then, the fabric was dipped in the solution and left in the water bath. 1g/50 ml Na2CO3 was added to the dyeing solution when the dyeing temperature was 60-70oC, and the sample was stirred for 45 minutes. The liquor ratio used for dyeing was 1:10.

#### 3% Dyeing Depth Shade

For 3% depth shade dyeing, 50 ml water, 0.15g Reactive dye, and 4g/50 ml NaCl were taken in a beaker. Then, the fabric was dipped in the solution and left in the water bath. 1g/50 ml Na2CO3 was added to the dyeing solution when the dyeing temperature was 60-70oC, and the sample was stirred for 45 minutes. The liquor ratio used for dyeing was 1:10.

#### 1% Dyeing Depth Shade

For 1% depth shade dyeing, 50 ml water, 0.05g Reactive dye, and 4g/50 ml NaCl were taken in a beaker. Then, the fabric dipped in the solution and left in the water bath. 1g/50 ml Na_2_CO_3_ was added to the dyeing solution when the dyeing temperature was 60-70°C, and the sample was stirred for 45 minutes. The liquor ratio used for dyeing was 1:10.

### Wash–off Process

After the dyeing process, a conventional wash-off was used to remove unfixed dyes from Standard-Dyed Fabrics. The conventional wash-off process includes many wash-off steps at different temperatures. Dyed fabrics were washed off according to the standard method of stirring fabric, following 1:10 in each step.

### Batch Wash-off Process

The Standard wash-off wastewater of each dye and shade depth was divided into 4 equal parts, each 335 ml, in a 1000 ml beaker. The remaining 3 Batch Fabrics of each dye and shade (1%, 3%, 5%) were washed off by treating the standard wash-off wastewater by Electrocoagulation.

### Electro-Coagulation lab setup description

The spent standard wash-off liquor was taken in a 1000ml beaker or graduated cylinder and then divided into three equal parts. pH was maintained at 4, 7, and 10 (acidic, neutral, and basic pH). Then, the electro-coagulator with two iron electrodes was inserted in each beaker for different periods (5min, 10min, 15min, 20min, 25min, 30min). After settling the sludge, the filtration assembly filtered the treated wash-off liquor. Then, this treated wash-off liquor was used for the wash-off process for batch samples.

### Wash Fastness

ISO 105; C06 method of wash fastness was used to determine the resistance of dyeing material on fabric after washing. The batch was washed, and values were visually evaluated by a grayscale, i.e., 1-5, representing shade change (Burkinshaw & Salihu, 2013).

## 3. Results & Discussion

### Effect of Electrolysis Time on Dye Removal

The effect of electrolysis time on EC performance was observed. A bench-scale laboratory procedure was conducted to measure the percentage of dye removal. The effect of electrolysis time illustrates the removal of reactive dyes as a function of operational parameters. Time is having a considerable impact on the removal of dyes.

### Effect of pH

Tables 3.2.1, 3.2.2, and 3.2.3 show the results of removing C.I. Reactive Dyes were significantly affected by a change in pH. To study this effect, the pH of standard wastewater was set to the desired values of 4, 7, and 10 by adding acetic acid (CH_3_COOH) and sodium hydroxide (NaOH).

### Effect of Electrolysis Time on Dye Removal

Electrolysis time influences treatment efficiency. Dye removal depends on the metal ions formed from the electrodes (Amour et al., 2016). An increase in electrolysis time leads to an increase in reaction time. (Zazou et al., 2019). This increase in time is attributed to the production of hydroxide ions (HO-), which make flocs and remove dyes from dyeing wastewater.

### C.I. Reactive Red 221

When the electrolysis time for 5% dye shade increased from 10 to 20 minutes, the removal efficiency increased from 55 to 93% and remained constant. The removal efficiency increased as dye concentration decreased. The removal efficiency of 3% was 96% in 20 minutes, i.e., more than the 5% dye shade. Excellent color removal was observed in 1% dye shade, i.e., 98% within 20 minutes at pH 10.

**Figure 1.**
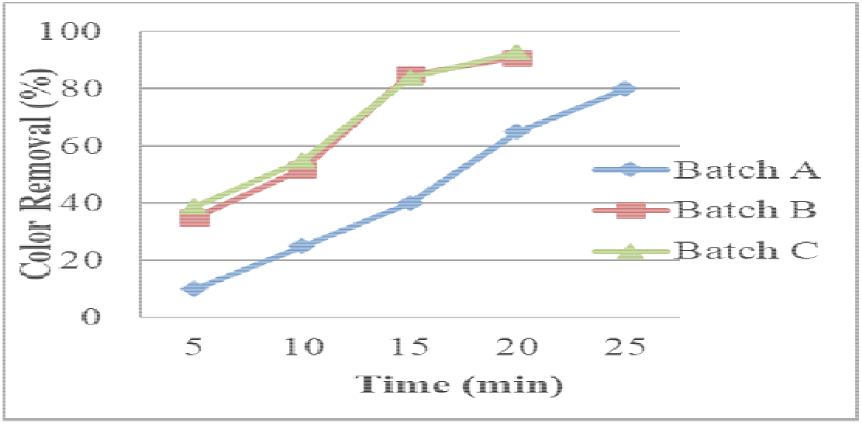
Color reduction percentage of C.I. Reactive Red 221 for 5% shade.

**Figure 3.**
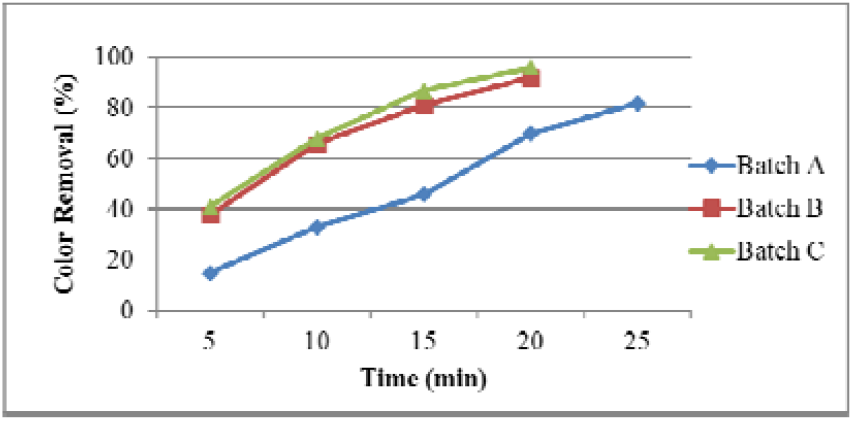
Color reduction percentage of C.I. Reactive Red 221 for 3% shade.

**Figure 4.**
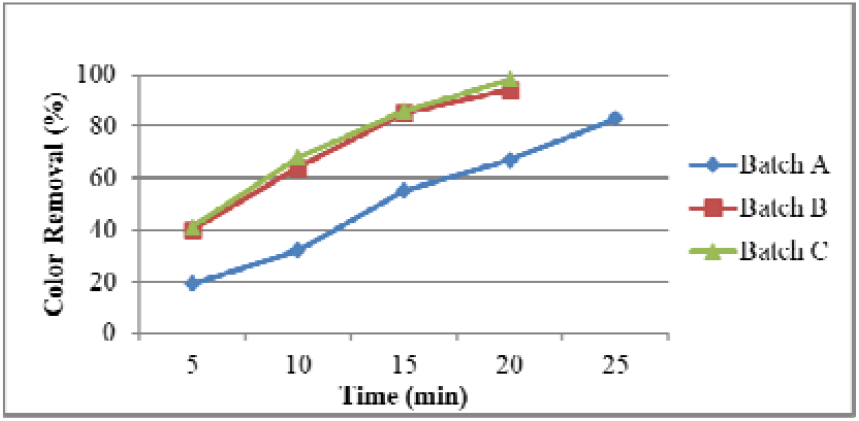
Color reduction percentage of C.I. Reactive Red 221 for 1% shade.

### C.I. Reactive Blue 19

The C.I. Reactive Blue-19 dye shades (5%, 3%, 1%) showed maximum color reduction within 10-20 minutes at pH 10, reaching over 96%. The last color reduction was observed at pH 4 with a maximum time range of 25 minutes, and further increased in time cannot enhance removal efficiency.

**Figure 5.**
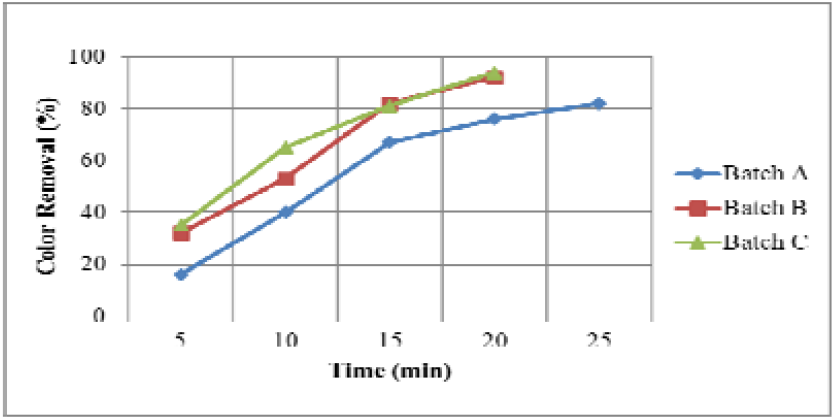
Color reduction percentage of C.I. Reactive Blue 19 for 5% shade.

**Figure 6.**
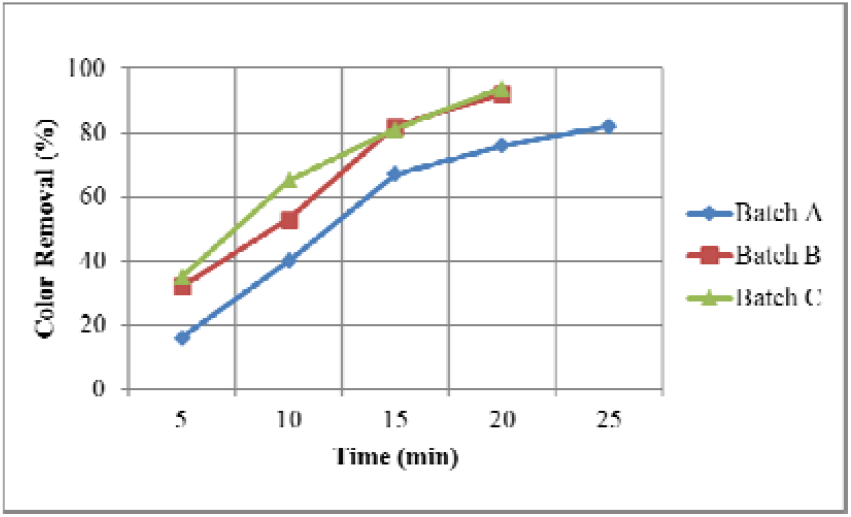
Color reduction percentage of C.I. Reactive Blue 19 for 3% shade.

**Figure 7.**
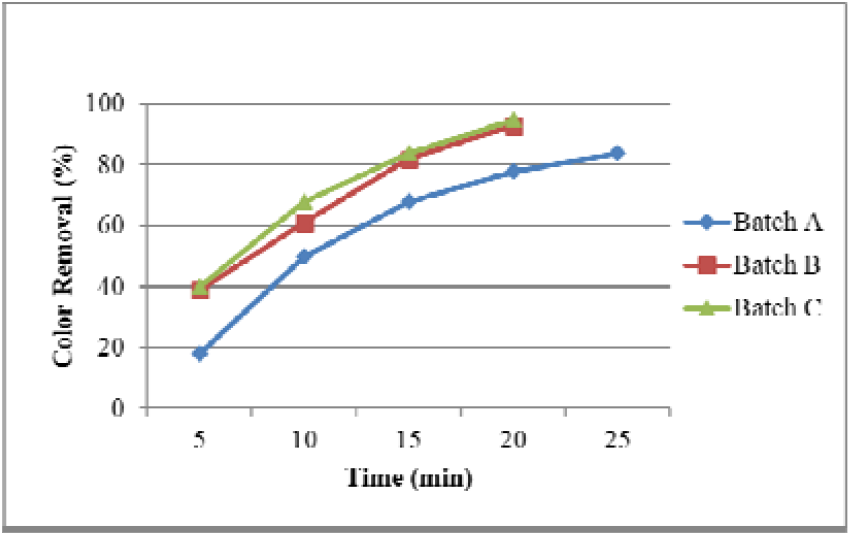
Color reduction percentage of C.I. Reactive Blue 19 for 1% shade.

### C.I. Reactive Yellow 145

The result of C.I. Reactive Yellow 145 with dye shades (5%, 3%, 1%) showed that as dye concentration decreases, the removal efficiency increases. pH 4 takes more time to remove dyes from dyeing wastewater than pH 7 and 10. The maximum removal efficiency of pH 4 was 84 %; after 25 minutes of further increased electrolysis time, the color reduction was unchanged.

**Figure 8.**
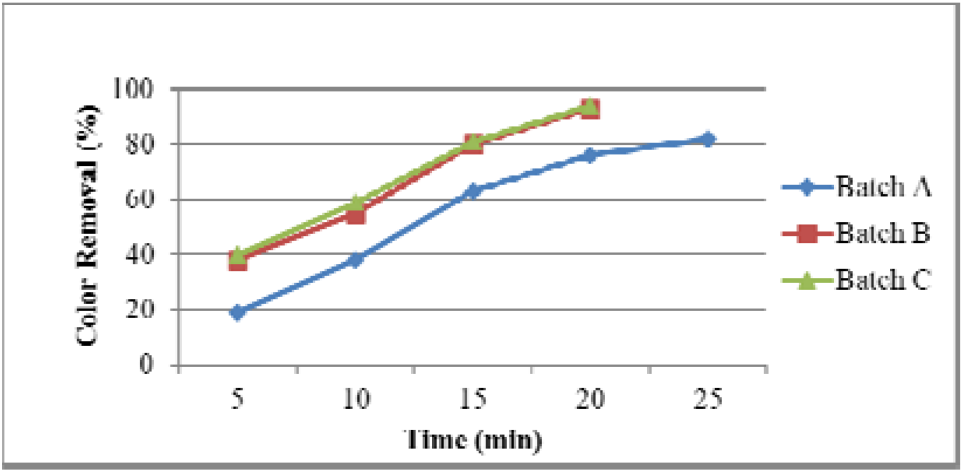
Color reduction percentage of C.I. Reactive Yellow 145 for 5% shade.

**Figure 9.**
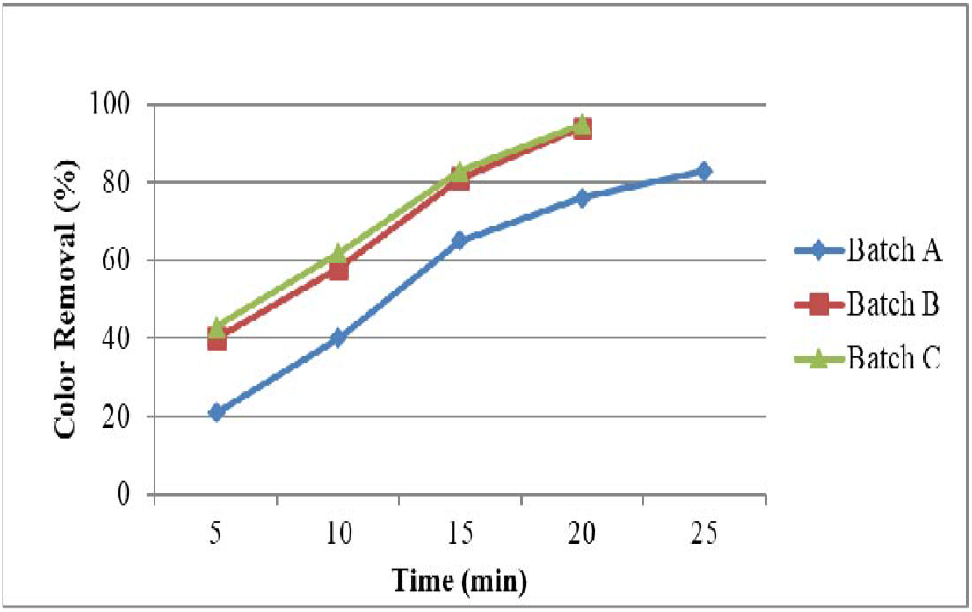
Color reduction percentage of C.I. Reactive Yellow 145 for 3% shade.

**Figure 10.**
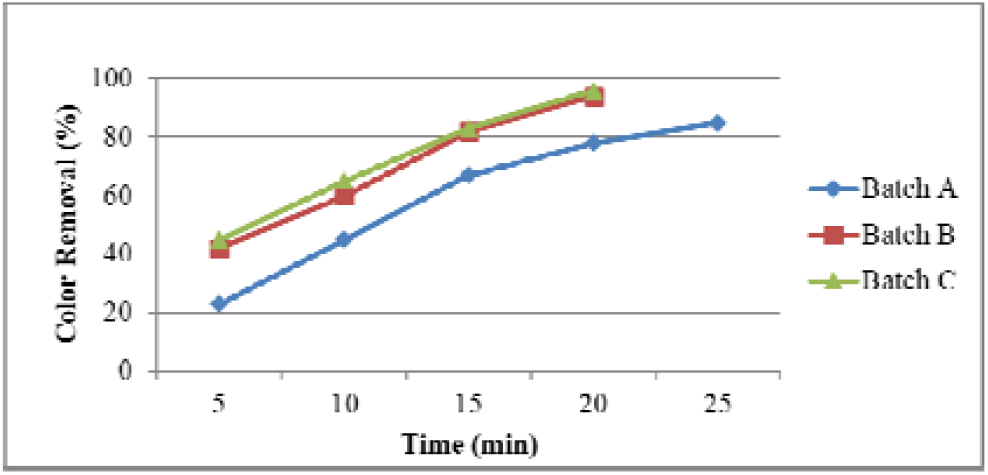
Color reduction percentage of C.I. Reactive Yellow 145 for 1% shade.

### Effect of pH

pH considerably influences the EC treatment method’s efficiency in removing dyes from dyeing wastewater. To study its effect, the pH of standard wastewater was set to the desired values of 4, 7, and 10 by adding acetic acid (CH_3_COOH) and sodium hydroxide (NaOH). The effect of pH 4 on EC efficiency indicates the poorest color removal, below 84%. However, excellent removal efficiency is achieved when pH reaches the maximum basic level. The experimental results showed that when pH was changed from 7 to 10 of dyeing wastewater, the removal efficiency increased.

**Figure 2.**
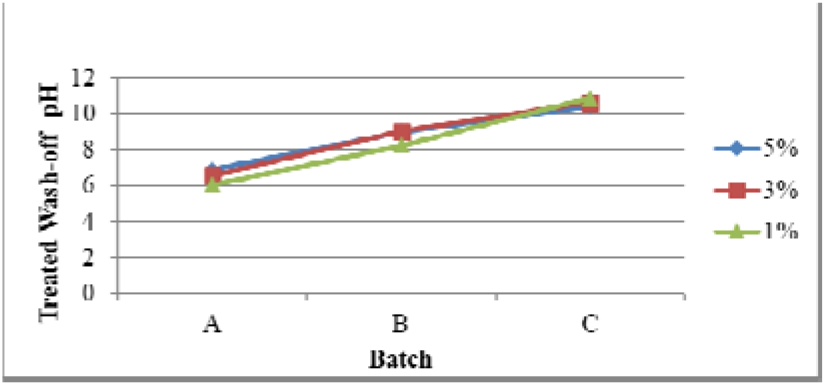
The pH of Treated wash-off of C.I. Reactive Red 221.

**Figure 3.**
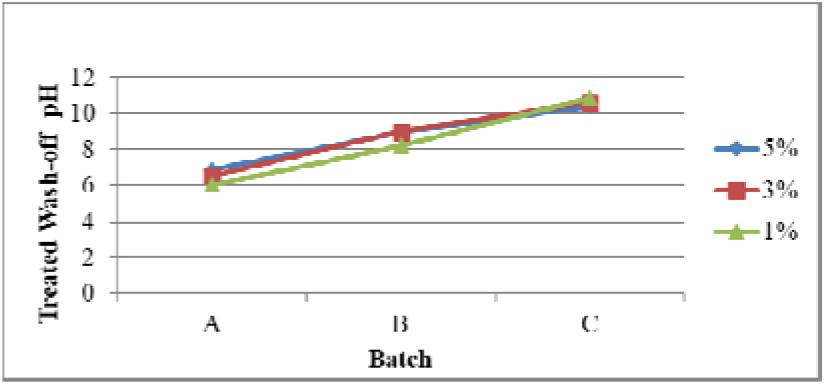
The pH of treated wash-off of C.I. Reactive Blue 19.

**Figure 4.**
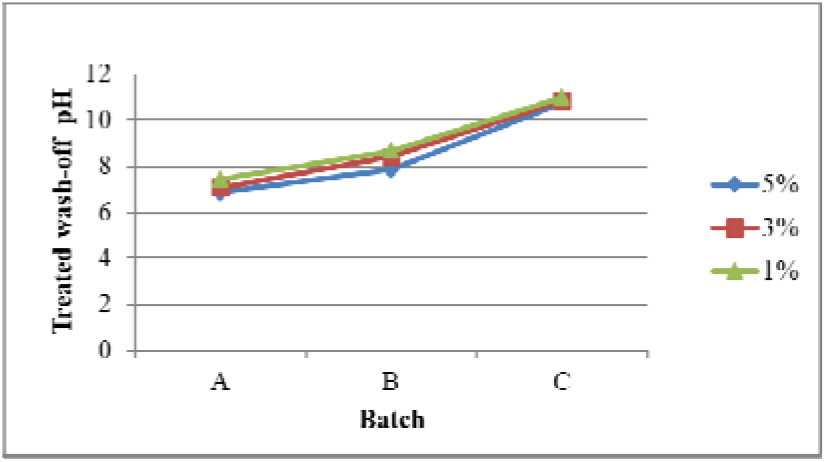
The pH of treated wash-off of C.I. Reactive Yellow 145.

### Wash Fastness

The ISO 105; C06 method of wash fastness was used to determine the resistance of dyeing material on fabric after washing. The batch was washed, and values were evaluated visually using grayscale, i.e., from 1 to 5, representing shade change.

| Gray Scale |  |
| --- | --- |
| Values | Shade Change |
| 1. | Much change |
| 2. | Considerable change |
| 3. | Noticeable change |
| 4. | Slightly change |
| 5. | Negligible change |

### Determination of wash fastness

ISO 105; C06\C2S and wash test using SDC multi-fiber was used to check the wash fastness of standard and batches. This method involves the washing of batches five times. After washing, the batches were assessed in the presence of grayscale by the ISO 105; A02 test protocol, and cross-staining wash-off was governed by ISO 105; A03 test protocol. The grey scale ranges from 1 to 5. 1 indicates maximum change in shade, while 5 represents no change in shade(Chantes et al., 2015).

## 4. Conclusions

The research on electrocoagulation as an effective process for treating textile wastewater has yielded promising results. The study’s main objective was to investigate the efficiency of color removal through electrocoagulation treatment and explore its potential for water reuse, addressing the issue of water scarcity. To achieve this, three different C.I. Reactive Dyes, namely Red 221, Yellow 145, and Blue 19, were used to dye cotton fabric, and the resulting wash-off was reused. The dye wastewater was treated at varying pH levels, such as 4, 7, and 10, using the electrocoagulation (EC) technique, which proved to be a reliable and environmentally friendly method for removing both organic and inorganic matter from wastewater, contributing to environmental protection. Compared to other treatment methods, the EC process exhibited lower sludge formation. Notably, a % removal efficiency of 96% was achieved for the C.I. Reactive Dyes, particularly at a pH of 10 within a time interval of 15-20 minutes. Evaluation of dry and wet crocking results indicated minimal unfixed dye, suggesting good wash fastness whose values ranged from 4 to 5. These results signify excellent color fastness, indicating little to no staining. Furthermore, the total color difference values for Batch C (fabric treated at pH 10) of the C.I. Reactive Dyes used in the study fell within acceptable limits (Burkinshaw & Salihu, 2013), validating the effectiveness of the electrocoagulation treatment process.

## Acknowledgments

I want to acknowledge and express my sincere gratitude to the authors for their dedicated work and valuable insights in crafting this study. I also extend my great appreciation to the laboratories involved, whose state-of-the-art facilities and expertise contributed to the accuracy and reliability of the results.

